# Confluent growth state dependent transcriptomic adaptation in A549 lung cancer cells

**DOI:** 10.64898/2026.08.27.747534

**Authors:** Akalya Sendrayakannan, Nidhi Yadav, Ashutosh Sahoo, Ranjan Kumar Nanda, Shyam Kumar Masakapalli

**Author notes:** Corresponding:* (*S.K.M*), (*R.K.N*).

## Abstract

Cell confluency is a major determinant of cell-cell communication, protein interactions, access to nutrients, and signalling dynamics, thereby significantly impacting biological outcomes. Lung cancer cells like A549 are widely used as screening models for scientific studies wherein their growth *in vitro* progress from non-confluent to confluent growth. In this study, we investigated the transcriptomic adaptations associated with the transition of A549 cells from baseline non-confluent to confluent growth. Comparative transcriptomic analysis between confluent and cells at baseline identified 815 upregulated and 671 downregulated transcripts. Pathway enrichment analysis of deregulated transcripts in confluent cells revealed enhanced cholesterol and sterol biosynthetic pathways, along with suppression of chromosomal segregation and mitotic pathways. At confluency, an increased expression of glucose transporters (SLC2, SLC60, and SL37 families) and glycolytic pathways, and a decrease in amino acid transporters (SLC1, SLC7, SLC38, and SLC36) and amino acid metabolic pathways is observed. A reduced one-carbon metabolic signature (SHMT2, DHFR, and MTHFD2) and enhanced fatty acid precursor synthesis (HMGCLL1, ALDH6A1, and AASS) were also observed at confluency. ^1^H NMR profiling of culture media revealed higher glucose and glutamine utilisation with lactate accumulation during culture maturation. Collectively, the data suggest transcriptome-level rewiring in A549 cells with preferential biosynthesis of lipids and sterols at confluency and underscore the importance of considering culture maturity in cancer biology, metabolism, and therapeutic studies.

**Highlights:**

- Transcriptome of confluent A549 lung cancer cells deciphered.
- A549 Cells at confluency show enhanced cholesterol and steroid metabolism with reduced cell division by inhibiting chromosomal segregation and separation.
- The transcriptome of A549 cells highlights a metabolic shift of reduced one-carbon metabolism and enhanced fatty acid precursor synthesis.

**Graphical abstract:** 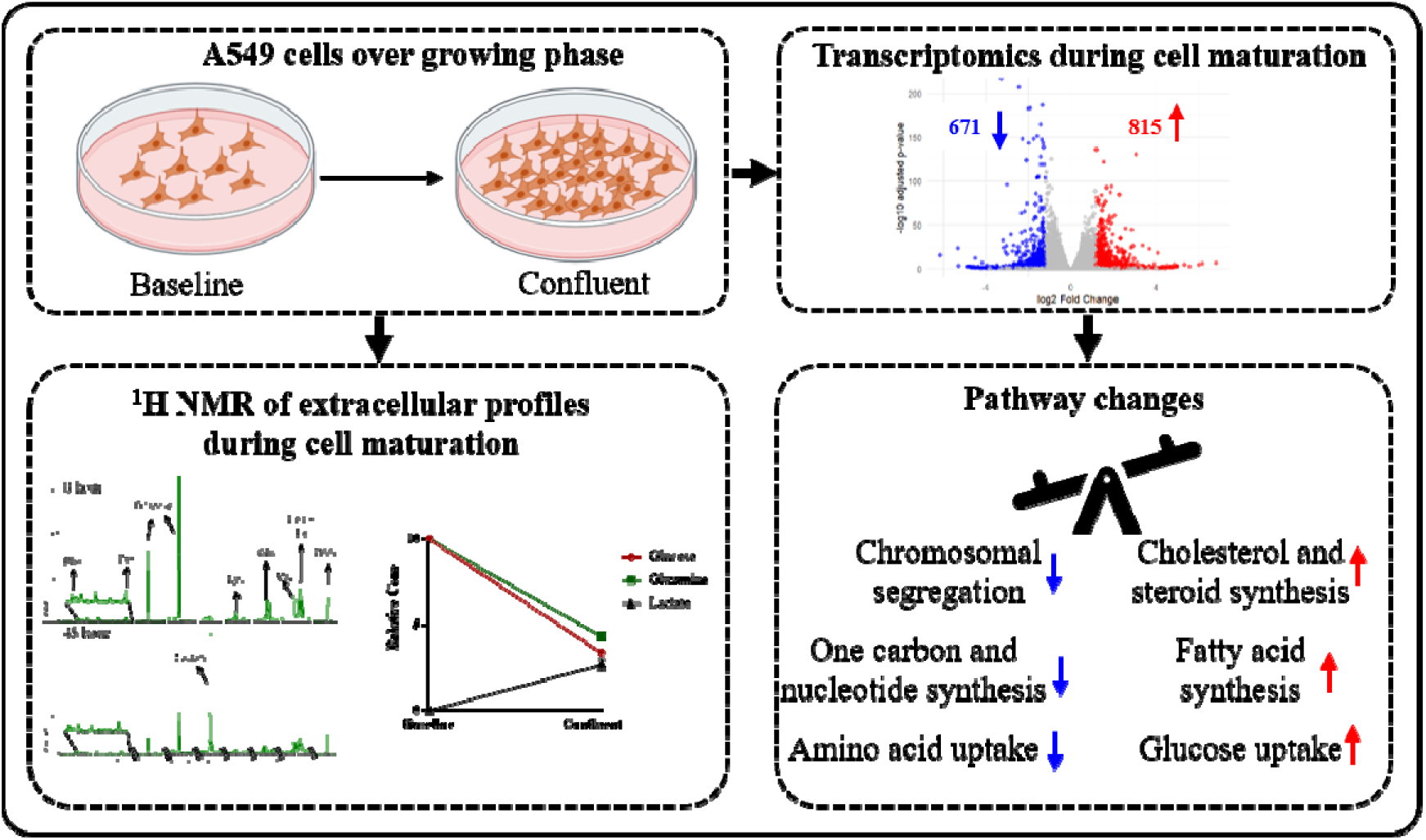

## 1. Introduction

Mammalian cell lines have long served as indispensable experimental model systems for studying cancer biology, therapeutic targets, and cellular responses under controlled conditions (Yu et al., 2019). Despite their widespread application in translational research, cell line models exhibit several discrepancies when compared to *in vivo* tumor systems. These differences largely arise from culture-associated metabolic alterations that distinguish them from the physiological human tumor microenvironment (Hynds et al., 2018). Among the various experimental factors influencing cellular behaviour, cell confluency is a major determinant, as it directly affects cell–cell communication, protein interactions, nutrient availability, and signalling dynamics, thereby significantly impacting biological outcomes (Xue et al., 2023).

The A549 lung adenocarcinoma cell line is one of the most widely used in vitro models in lung cancer biology, host-pathogen interaction studies, pulmonary research, and respiratory virology, particularly for investigating respiratory syncytial virus (RSV) infection (Saleh et al., 2020). Derived from human alveolar epithelium, A549 cells exhibit several characteristics of alveolar type II (AT II) epithelial cells, and prolonged cultivation has been reported to enhance their AT II-like phenotypic features (Cooper et al., 2016). Since drug discovery and pharmacological studies in mammalian cell lines are commonly performed at approximately 70–80% confluency (Gao et al., 2019), understanding the system-level transcriptomic adaptations and metabolic changes that occur during the transition from actively proliferating to confluent growth states is of considerable biological and translational importance.

Cancer cells exhibit remarkable metabolic plasticity and continuously modulate extracellular nutrient uptake and metabolite secretion in response to alterations in growth state, nutrient availability, and cellular density. Therefore, assessing extracellular metabolite utilization can provide functional insights into the metabolic state of cells and their adaptive responses during in vitro growth (Jain et al., 2012). In this context, nuclear magnetic resonance (NMR)- based metabolic profiling serves as a robust approach for monitoring extracellular metabolite consumption and release patterns, thereby enabling a more comprehensive understanding of cellular metabolic adaptation.

Hence, the present study investigates nutrient uptake and transcriptomic alterations in A549 cells during *in vitro* growth, with particular emphasis on amino acid and central carbon metabolism, as well as transporter-mediated nutrient exchange. By integrating transcriptomic profiling with NMR-based extracellular metabolite analysis, this study aims to elucidate the system-wide cellular adaptation at the transcript level and nutrient utilization dynamics in A549 lung cancer cells.

## 2. Materials and methods

### 2.1. Cell culture

A549, a type II alveolar epithelial adenocarcinoma cell line, was procured from NCCS, Pune. Cells were maintained and cultured in DMEM media (D9802-01, US Biological) with 25 mM glucose (G7021, Sigma), 4 mM glutamine (G8540, Sigma), 10% Dialyzed FBS (26400-044, Gibco), 0.4 mM serine (S4500, Sigma), and glycine (G7126, Sigma) Sodium pyruvate (TCL015, Himedia), Sodium bicarbonate (S5761, Sigma), and 1% (v/v) Penicillin-streptomycin (15140148, Gibco) in a humidified atmosphere at 37° C with 5% CO_2_.

### 2.2. Carbon source utilization from cell culture filtrate estimation

Cells were seeded at a density of 0.32 - 0.35 million per well in 6-well plates for the initial 32 hours. The media change or replenishment was done after the cells were gently rinsed with PBS. The spent media is harvested at the final time point, and both spent and unused media are lyophilized and stored for further analysis (Figure 1). Analysis was performed using ^1^H NMR spectroscopy (JEOL-Resonance-JNM-ECX-500 MHz) with D_2_O (151882, Sigma) as the solvent and DSS (3-(Trimethylsilyl)-1-propanesulfonic acid-d6 sodium salt; 178837, Sigma) as the internal standard. The parameters used were 64 scans with a relaxation delay of 5 seconds, a pulse width of 5.375 µs (45 ° angle), and a spectral width of 12 ppm. The spectra were processed manually using JEOL-Delta software (version 5.3.1), normalized by setting the DSS area to 1, and then further normalized to account for the protein content of the cells. Further, the initial concentration, as indicated by the manufacturer’s information, was used to calculate the absolute consumption. The plots were generated using GraphPad Prism, and Welch’s t-test was used for comparing the conditions.

**Figure 1.**
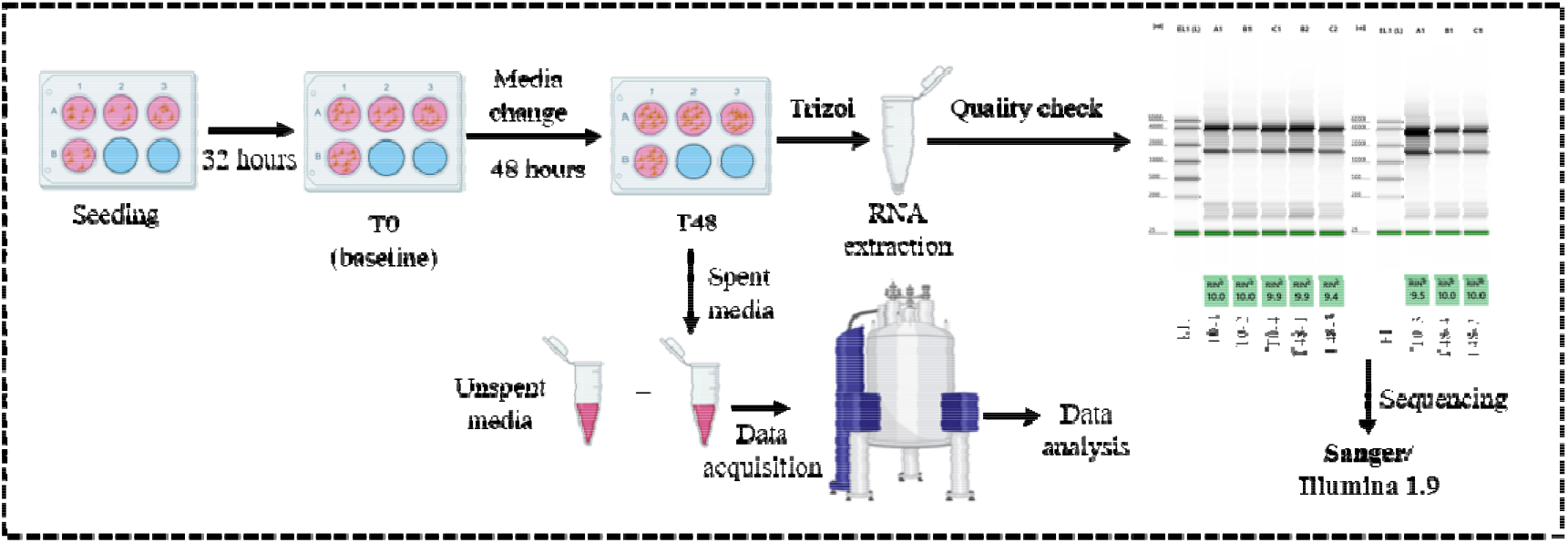
Overview of the workflow followed for harvesting samples for extracellular nutrient utilisation and RNA extraction.

### 2.3. RNA extraction and sequencing

The total RNA was extracted at baseline (0 hour-before media change) and 48 hours after the media change using TRIzol reagent according to the manufacturer’s instructions (Invitrogen). Briefly, 0.2 mL of chloroform was added to each sample, which was then incubated for 2-3 minutes, followed by centrifugation at 12000 × g at 4 °C for 15 minutes. The aqueous phase was taken, and 0.5 mL of propanol was added. The mixture was then incubated at 4 °C for 10 minutes, followed by centrifugation at 12,000 × g for 10 minutes at 4 °C. The pellet was collected, and 1 mL of 75% ethanol was added. The mixture was vortexed briefly and then centrifuged at 7500 × g for 5 minutes at 4 °C. The pellet was air-dried, resuspended in RNase-free water, and stored at -80 °C until further use. The RNA was quantified using a Qubit fluorometer with a broad-range (BR) assay kit (Q10210, Invitrogen), and the RNA integrity (RIN) was assessed using TapeStation (Agilent) with a broad-range RNA kit. The sequencing was performed on an Illumina Novaseq 6000 using paired-end reads by NGB Diagnostics Private. Limited., New Delhi.

### 2.4. Transcriptomics data analysis

The raw RNA reads were checked for quality using fastQC, a package from the SRA toolkit (sratoolkit.current-ubuntu64.tar.gz, Leinonen et al., 2010). The reads were then aligned to the indexed human reference genome (GRCh38.p14), using STAR (https://github.com/alexdobin/STAR, Dobin et al., 2013). The coordinated sorted files were then counted for features using feature count, a package in subreads (Liao et al., 2014). These were carried out in the high-performance computing facility of IIT Mandi, Himachal Pradesh, India with an in-house pipeline. The counted files were then subjected to DESeq2 for the differential expression analysis in R (Love et al., 2014). The transcripts were annotated using the GO database and org.Hs.eg database (https://bioconductor.org/packages/org.Hs.eg.db/ and Manjang et al., 2020). The genes with log_2_ FC ≥ ±1.2 and adjusted P-value < 0.05 were considered deregulated and included for further analysis of over-representation analysis for pathway decoding (Cluster profiler, Yu et al., 2012). The volcano plots were generated using the ggplots package of R (Wickham, 2016). The heatmaps were generated (log_2_ FC ≥ ± 0.5 and adjusted P-value of 0.05) by normalizing each gene’s expression to z-scores using the heatmaply package in R (Galili, 2017). The genes for the pathway analysis were collected from the KEGG database (Kanehisa & Goto, 2000).

## 3. Results and Discussion

### 3.1. Macromolecule utilization of cells over the growth period

The A549 cells were studied at confluent stage, as shown in Figure 1. The major macromolecules, including sugars and amino acids, were studied by NMR, as shown in Figure 2a. The relative percentage of glucose remaining in the media during the confluency stage was decreased to 25% during the confluent phase (Figure 2b). The lactate released into the media was increased at 2.67 DSS relative levels (Figure 2h). Glutamine, the second-most consumed macromolecule after glucose, was also consumed rapidly, and the relative percentages remaining in the media was 32% at the end of 48 hours (Figure 2c). Other amino acids were also measured, as mentioned (Figure 2d, 2e, 2f, 2g). However, their consumption was not beyond 50%. These macromolecular consumptions and excretion were closely associated with the previous studies (O’Neill et al., 2022). Further, it indicated that the major macronutrients were not being starved in the cells, raising questions about their metabolism and the expression of the transporter responsible for their uptake. Further, the cells’ metabolic pathway shift at confluency was of interest.

**Figure 2.**
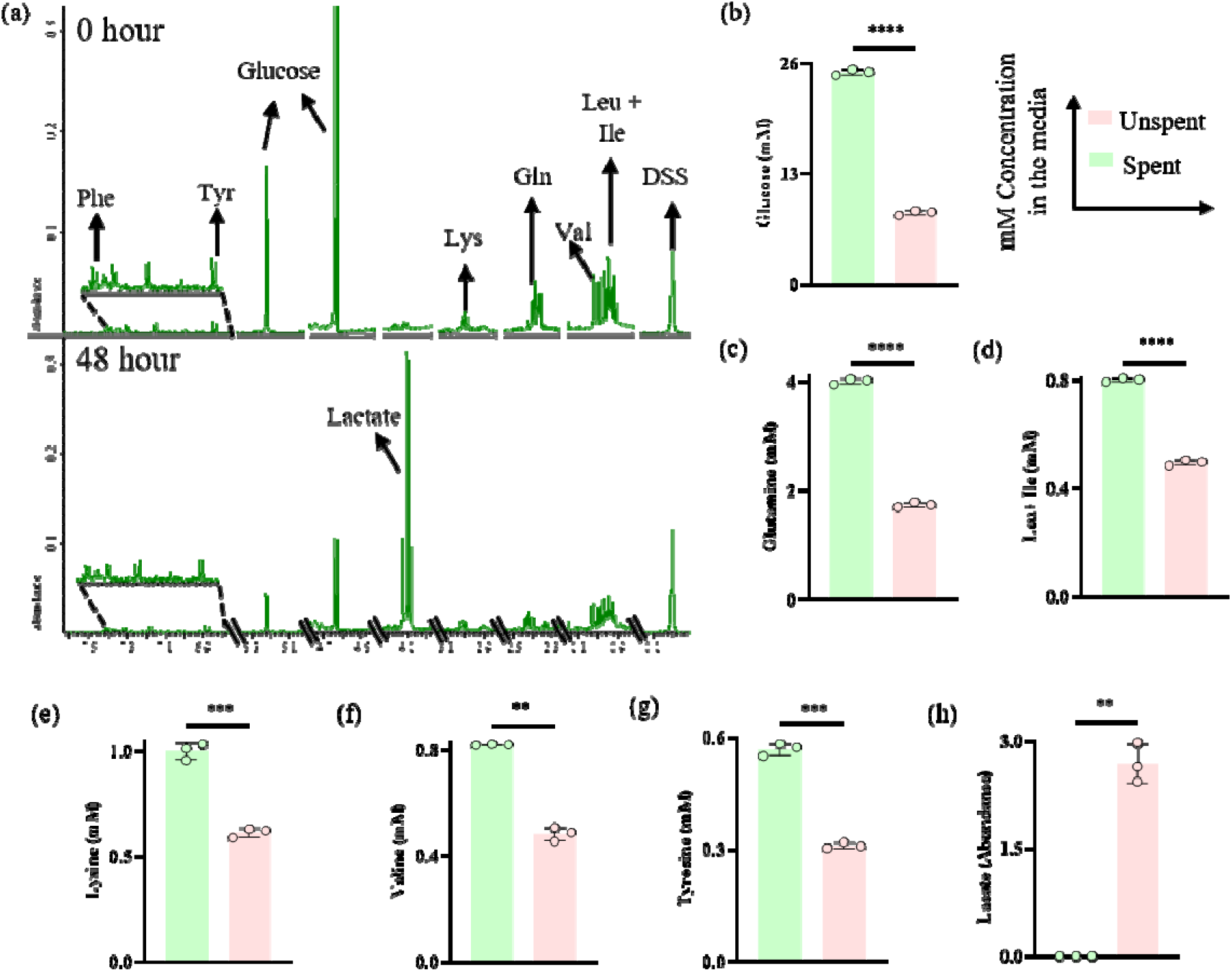
Nutrient utilisation and lactate production by A549 cells. (a)The levels of nutrients in DMEM media (spent and unspent) analysed by 1H NMR using DSS as an internal standard. The concentration of (b) glucose, (c) Glutamine, (d) Leucine + Isoleucine, (e) Lysine, (f) Valine, (g) Tyrosine, and (h) lactate in the spent and unspent media. Lactate is presented as a relative abundance. **p≤ 0.005, ***p≤ 0.0005, ****p≤ 0.00005 (Welch’s t-test).

### 3.2. Overall transcriptomics shows changes associated with the maturation of A549 cells

The quality of all sequenced RNA samples was robust, with RNA Integrity Number (RIN) scores ranging from 9.4 to 10, indicating negligible RNA degradation (Supplementary Table S1). After sequencing, the FastQC files were subjected to the pipeline as indicated in Figure 3a. Sequencing performance was high across all libraries, showing optimal results for base quality, adapter content, and GC distribution as confirmed by FastQC analysis. Alignment to the human reference genome using the STAR tool was highly efficient, yielding 85–89% of reads uniquely mapped. The remaining read fractions consisted of multi-mapping reads (7.5– 8.7%) and reads excluded for being too short (2.9–8.8%). Feature quantification was performed via featureCounts using stringent criteria (flags -B and -C) to count only complete, non-ambiguous pairs. Under these conservative parameters, the assignment rate remained consistent at 60–67%, confirming the high quality of the transcriptomic data and suitability for downstream differential expression analysis.

**Figure 3.**
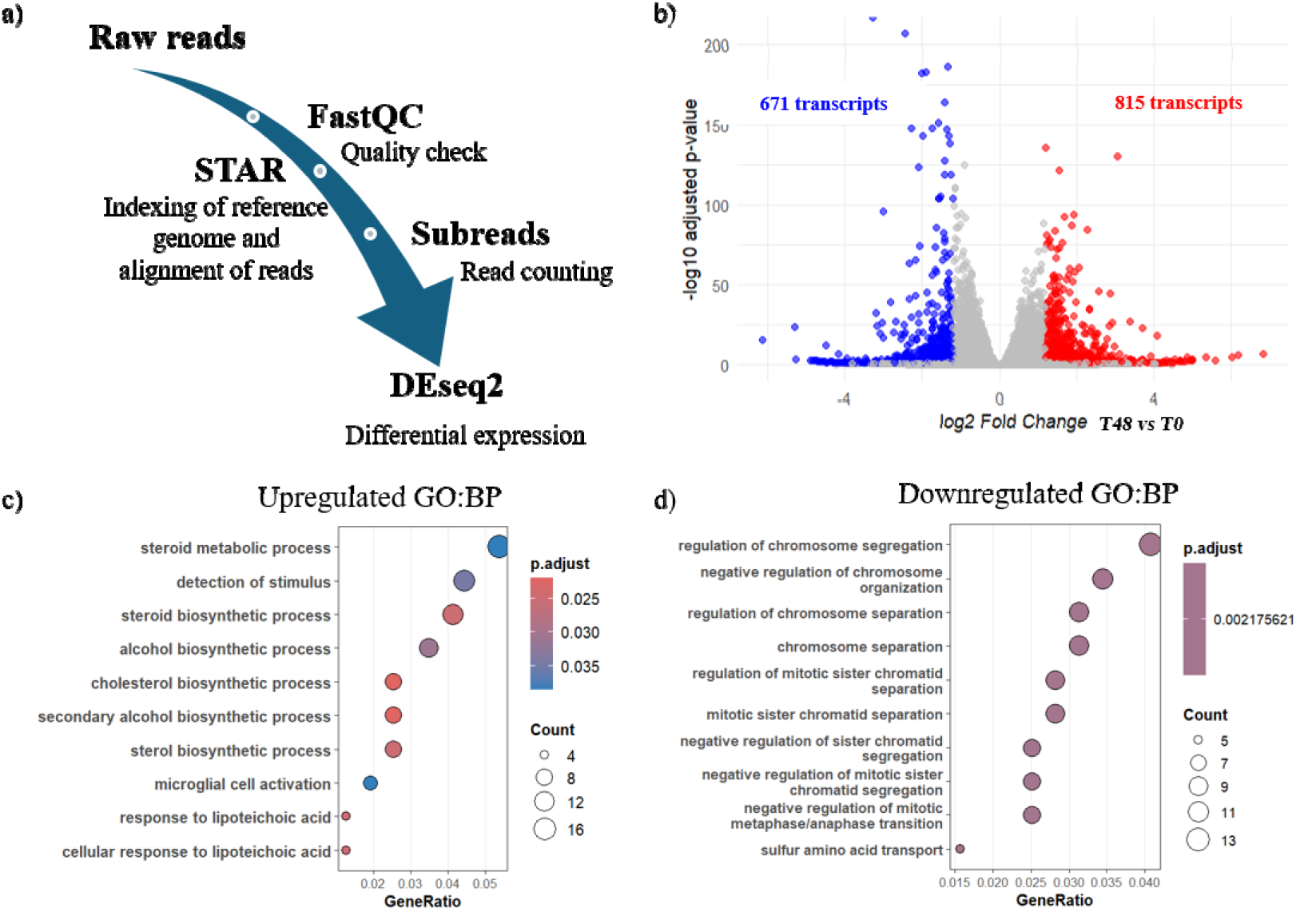
Confluency-associated transcriptomic changes of A549 cells. (a) Bioinformatics pipeline followed to obtain differential expression of transcripts. (b) Volcano plot showing the significance of alterations in transcripts between the basal time point (T0) and 48 hours (T48). The threshold for deregulated transcripts is a log2 fold change (T48 vs T0) of ± 1.2 and an FDR (adjusted p-value) of 0.05. Overrepresentation analysis with the GO (Gene Ontology) database for Biological Process (BP) showing the (c) upregulated and (d) downregulated pathways with cell maturation and confluency. The colour and size of the dot represent the significance and number of transcripts in the clusters. The Benjamin Hochberg method was used to control the FDR.

Further, with cell maturity, the transcript changes were clearly evident, as evidenced by the deregulated transcripts. There were 815 upregulated and 671 downregulated transcripts at 48 hours compared to the basal time point (Figure 3b). Further, over-representation analyses were performed using GO.db to identify upregulated and downregulated pathways associated with cell maturity. The pathways that are upregulated (Figure 3c) include the cholesterol and sterol biosynthetic process. The downregulated pathway (Figure 3d) is primarily associated with chromosomal segregation and organization.

### 3.3. Pathway alterations associated with transcript changes of maturing A549 cells

#### 3.3.1. Upregulation of sterol metabolism

The cholesterol and sterol biosynthetic process clusters around the transcripts HSD17B7, MSMO1, HMGCS1, HMGCR, INSIG1, FDFT1, IDI1, and APOE (Figure 4a and 4b). Further information on the clusters and statistics is provided in the Supplementary Table S2. All these enzymes involved in squalene, cholesterol, and steroid synthesis have been shown to contribute to invasion in lung cancer cells (Yang et al., 2020). The mevalonate pathway transcripts HMGCS1 and HMGCR are predominantly upregulated in cancers and may contribute to progression, invasion, and metastasis (Ashida et al., 2017). The response to lipoteichoic acid is mediated by TLR2, CD14, CD36, and LBP, and may be driven primarily by cell crowding or confluency, thereby preparing the system to elicit an immune response. Further, TLR overexpression is also indicative of proliferation and anti-apoptotic activity (Sato et al., 2009).

**Figure 4.**
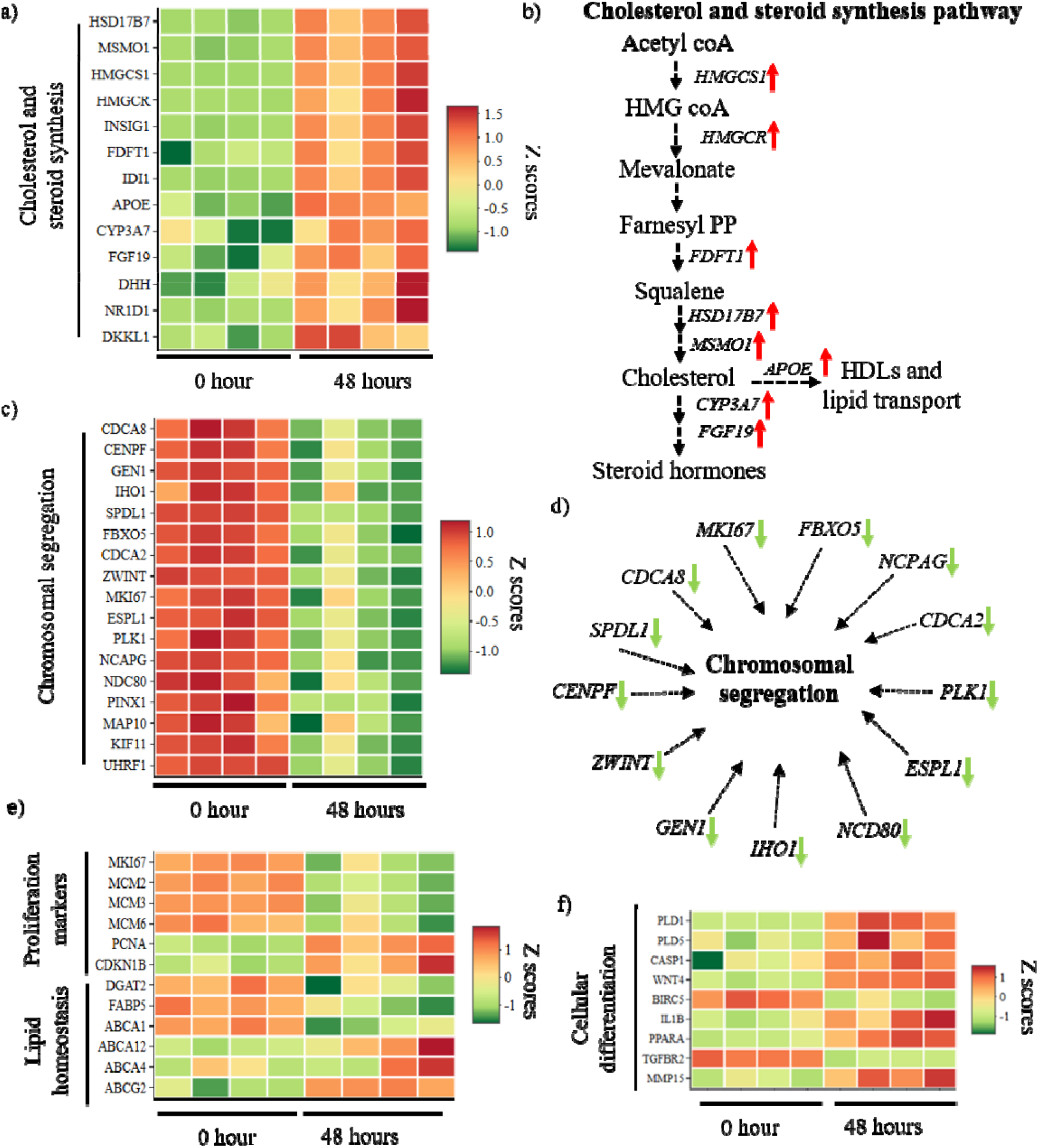
Transcriptomic alterations in pathways during A549 cell confluency. Row-normalized heatmaps (Z scores) of transcripts associated with (a and b) cholesterol and steroid metabolism, and its pathway model, (c and d) chromosomal segregation, and its model, (e) expression of proliferation and lipid homeostasis, and (f) cellular differentiation in A549 cells during maturation (T0 and T48). A variance-stabilizing transformation (VSD) was used to normalize transcript expression values (n=4). The transcripts were identified using a log2 fold change threshold of ±0.5 and an FDR (adjusted p-value) of 0.05, and deregulation was indicated by the color bars to the right of the heatmaps, with red indicating upregulation and green indicating downregulation. HMGCS1-3-hydroxy-3-methylglutaryl-CoA synthase 1; HMGCR-3-hydroxy-3-methylglutaryl-CoA reductase; FDFT1-Farnesyl-Diphosphate Farnesyltransferase 1; HSD17B7-Hydroxysteroid 17-Beta Dehydrogenase 7; MSMO1-Methylsterol Monooxygenase 1; APOE-Apolipoprotein E; FGF19-Fibroblast Growth Factor 19; CDCA8-Cell Division Cycle Associated 8; CENPF-Centromere Protein F; MKI67-Ki-67 protein; FBXO5 (F-box only protein 5); NCAPG gene-Non-SMC Condensin I Complex Subunit G; ZWINT (ZW10 Interacting Kinetochore Protein; SPDL1-Spindle Apparatus Coiled-Coil Protein 1; GEN1-Holliday junction resolvase; NCD80-kinetochore complex component; IHO1-Interactor Of HORMAD1 1; MCM (Minichromosome Maintenance; PCNA (Proliferating Cell Nuclear Antigen; CDKN1B (Cyclin Dependent Kinase Inhibitor 1B; DGAT2-Diacylglycerol O-Acyltransferase 2; FABP5 (Fatty Acid Binding Protein 5; ABCA-ATP-binding cassette (ABC); PLD1-Phospholipase D1; CASP1-Casapase-1; WNT4 (Wnt Family Member 4); BIRC5-Baculoviral IAP Repeat Containing 5; TGFBR2 (Transforming Growth Factor Beta Receptor 2; IL1B-Interleukin-1 beta; PPARA-Peroxisome Proliferator-Activated Receptor Alpha; MMP15-Matrix Metallopeptidase 15.

#### 3.3.2. Downregulation of chromosomal segregation

The process of chromosomal segregation and organization falls under the clusters that include the transcripts CDCA8, CENPF, GEN1, IHO1, NCAPG, SPDL1, FBXO5, CDCA2, ZWINT, MKI67, ESPL1, PLK1, and NDC80 (Supplementary Table S2). However, most cell division-associated (CDCA) transcripts are actively expressed in cancer cells to support cell division (Liu et al., 2024). These transcripts are downregulated as cells enter confluency; they spend less energy on cell division and more on cholesterol and steroid biosynthesis, thereby suggesting a confluency-associated quiescence (Figure 4c and 4d).

#### 3.3.3. Downregulation of proliferation markers and upregulation of ATP lipid transporters

The markers of AT II cells and their differentiation were studied based on the earlier study by Cooper et al. (2016), as shown in Figures 4e and 4f. Markers of alveolar type II epithelial cells, such as surfactant proteins, were studied, and no significant changes over time were observed. Also, stem cell and differentiation markers, as well as complement component markers, were studied, but none were significantly altered over time. The ATP lipid transporters ABCA12, ABCB4, and ABCG2 were upregulated; however, ABCA1 was downregulated, suggesting that lipid metabolism homeostasis in the cells increased over time, consistent with the AT II phenotype. The ATP lipid transporters have also changed to maintain lipid transport and homeostasis, as evidenced by the downregulation of ABCA1, which indicates reduced cholesterol efflux (He et al., 2020), and the upregulation of other transporters involved in ceramide transport to the cell wall (Akiyama, 2014), and upregulation of multidrug resistance enhancement through upregulating ABCG2 and ABCB4 (To et al., 2011; Kiehl et al., 2014). Furthermore, the downregulation of FABP5 indicates its reliance on AT II characteristics (Yin et al., 2026). Proliferation markers, such as MKI67 and MCM, were downregulated, indicating decreased proliferation upon confluency; however, the PCNA transcript was upregulated, suggesting activation of the repair mechanism at the end of the cell cycle (Juríková et al., 2016). The cell cycle inhibitor CDKN1B is also upregulated, further indicating that the cells are not actively dividing (Haferlach et al., 2011). Furthermore, as cell cycle inhibition is directly associated with lipid droplet formation and DGAT2 regulation, its downregulation would be directly related to cell cycle arrest (Liu et al., 2024a).

Cellular differentiation was studied as shown in Figure 4f, and it was observed that, over time, A549 cells activated Wnt4, MMP15, PLD, CASP1, and IL1B, while downregulating BIRC5 and TGFBR2. The PLD upregulation directly correlated with Wnt4 upregulation, which is directly associated with tumor progression and epithelial-to-mesenchymal transition (Lim et al., 2024). In addition, MMP15 is upregulated and directly associated with epithelial-to-mesenchymal transition (EMT) (Khalili-Tanha et al., 2025). Further upregulation of Casp1, along with downregulation of BIRC5, indicates that the cell is entering the senescence pathway (Kahm & Kim, 2023). The overall mechanism, as evidenced by this, is the transition of cells from epithelial to mesenchymal states, which confers migratory and metastatic properties.

### 3.4. Understanding the transporters of macromolecules and their metabolism upon confluency of A549 cells

The macromolecules, including sugars, amino acids, and fatty acids, as well as their transporters and metabolism, were studied. The transporters under the families SLC1, SLC3, SLC6, SLC7, SLC15, SLC16, SLC17, SLC18, SLC32, SLC36, SLC38, SLC43, SLC66, and some of the transporters under SLC25 were studied for amino acid transporters. The transporters under the families SLC2, SLC5, SLC35, SLC37, SLC45, SLC50, and SLC60 were studied for sugar transporters. The transporters under the families SLC27, SLC33, SLC59, SLC63, and SLC65 were studied for fatty acid and lipid transport. The amino acid transporters and sugar transporters were inversely expressed, as seen in Figure 5a and 5c. The amino acid transporters were expressed at higher levels in the initial phase than in the confluency phase, except for the lysine and arginine transporters (SLC7A2 and SLC25A29), and the sugar transporters were expressed at higher levels in the confluent phase. Further, as seen from the metabolism of these compounds, it is directly correlated with the transporters (Figure 5b and 5c). Transcripts encoding for sugar metabolism in the central carbon pathway are higher at later time points than in the early phase, as seen in ALDOC, TKT, OGDHL, SDHD, and ACO2. The synthesis of some non-essential amino acids is upregulated, including arginine (ASS1), cysteine (CTH), and glutamate (GOT1). On the other hand, the non-essential amino acid synthesis pathways, including serine (PHGDH and PSAT1), proline (PYCR1), and alanine (GPT2), are downregulated.

**Figure 5.**
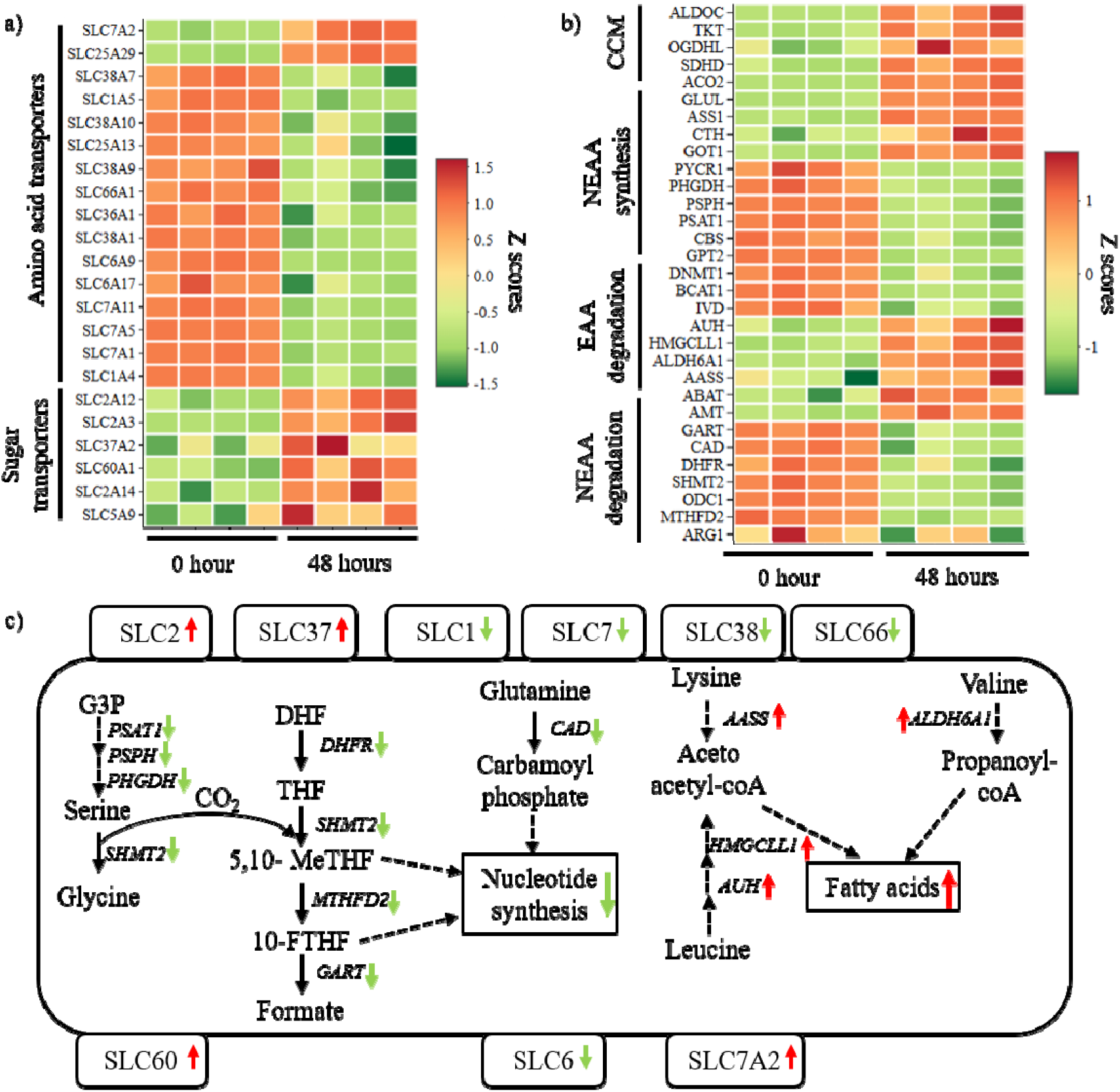
Transcript changes associated with nutrient uptake and utilisation in A549 cells upon confluency. a) Row-normalised heatmap (Z score) showing variations in the amino acid and sugar uptake upon growth of A549 cells. b) Heatmap showing variations in the metabolism of amino acid and sugar upon growth of A549 cells. c) Overall model for nutrient uptake and utilisation happening during cell confluency (T0 and T48). A variance-stabilizing transformation (VSD) was used to normalize transcript expression values (n=4). The transcripts were identified using a log2 fold change threshold of ±0.5 and an FDR (adjusted p-value) of 0.05, and deregulation was indicated by the color bars to the right of the heatmaps, with red indicating upregulation and green indicating downregulation.SLC-Solute carrier proteins (SLC25-mitochondrial transporter); CCM-Central carbon metabolism; NEAA-Non-essential amino acid; EAA-Essential amino acid; ALDOC-Aldolase fructose bisphosphate C; TKT-Transketolase; OGDHL-Oxoglutarate Dehydrogenase Like; SDHD-Succinate Dehydrogenase Complex Subunit D; ACO2-Aconitase 2; GLUL-Glutamate-Ammonia Ligase; ASS1-Arginino succinate Synthase 1; CTH-Cystathionine Gamma-Lyase; GOT1-Glutamic-Oxaloacetic Transaminase 1; PYCR1-Pyrroline-5-Carboxylate Reductase 1; PHGDH-Phosphoglycerate Dehydrogenase; PSPH-Phosphoserine Phosphatase; PSAT1-Phosphoserine Aminotransferase 1; CBS-Cystathionine Beta-Synthase; GPT2-Glutamic--Pyruvic Transaminase 2; DNMT1-DNA Methyltransferase 1; BCAT1-Branched Chain Amino Acid Transaminase 1; IVD-Isovaleryl-CoA Dehydrogenase; AUH-AU RNA Binding Methylglutaconyl-CoA Hydratase; HMGCLL1-3-Hydroxymethyl-3-Methylglutaryl-CoA Lyase Like 1; ALDH6A1-Aldehyde Dehydrogenase 6 Family Member A1; AASS-Aminoadipate-Semialdehyde Synthase; ABAT-4-Aminobutyrate Aminotransferase; AMT-Amino methyl transferase; GART-Phospho ribosyl glycinamide Formyl transferase; CAD-Carbamoyl-phosphate synthase 2; DHFR-Dihydrofolate Reductase; SHMT2-Serine Hydroxy methyl transferase 2; ODC1-Ornithine Decarboxylase-1; MTHFD2-Methylene tetrahydrofolate Dehydrogenase 2; ARG1-Arginase 1. Other transcripts studied related to these pathways are mentioned in the Supplementary Table S3.

The amino acid degradation pathway is active during the confluent phase (Figure 5b), but most amino acid pathway genes that support nucleotide synthesis via one-carbon metabolism are downregulated. The transcripts, including GART, DHFR, SHMT2, and MTHFD2, that are involved in the degradation pathway of non-essential amino acids, which are a part of one-carbon metabolism, are downregulated in the confluent phase. Additionally, the CAD, the primary transcript for nucleotide synthesis from glutamate, is downregulated (Cooper et al., 2016). The transcripts ARG1 and ODC1, which are involved in the urea cycle, are downregulated. Also, the transcripts AUH, AASS, HMGCLL1, and ALDH6A1, which are primarily involved in the final reactions of amino acids that produce acetoacetyl-CoA or propionyl-CoA, which are involved in fatty acid metabolism, are upregulated. This further confirms that cells at confluence are trying to maintain fatty acid synthesis by diverting from amino acids and nucleotide synthesis.

## Conclusion

This study provides transcriptomic adaptations that occur as A549 cells progress from an actively growing state to confluency. As the cells reach the confluent state, they utilise higher glucose and glutamine and lower consumption of other amino acids including valine, leucine, isoleucine, phenylalanine, lysine, and tyrosine.

Transcriptomic analysis revealed extensive growth-state-dependent remodelling, with 815 transcripts upregulated and 671 downregulated at confluency. The most prominent transcriptional changes included activation of cholesterol and sterol biosynthetic pathways and suppression of chromosomal segregation, mitotic progression, and proliferation-associated programs. The downregulation of canonical proliferation markers together with the upregulation of cell-cycle inhibitory mechanisms suggests a transition toward a non-proliferative state upon confluency. During confluency, the A549 cells ability to express the markers like surfactant proteins is not altered; however, the ABC transporters are upregulated.

Further investigation of nutrient transport systems revealed coordinated metabolic rewiring. Confluent cells exhibited increased expression of glucose transporters and central carbon metabolic genes, whereas amino acid transporters and pathways associated with amino acid biosynthesis and one-carbon metabolism were generally reduced, further indicating reduced nucleotide generation. Concurrently, pathways involved in amino acid catabolism and fatty acid precursor generation were enhanced.

Collectively, these findings demonstrate that confluency induces a coordinated shift in A549 cells from a highly proliferative, nucleotide-supporting metabolic program toward a lipid- and sterol-oriented metabolic phenotype. This transition is characterized by reduced cell-cycle activity, altered expression of nutrient transporters, suppression of one-carbon metabolism, and increased lipid biosynthetic capacity. The study highlights the substantial influence of growth state on cellular metabolism and gene expression and underscores the importance of considering culture maturity when using A549 cells as experimental models in cancer biology, metabolism, and therapeutic studies.

## Acknowledgement

Akalya Sendrayakannan (ASK) and Ashutosh Sahoo (AS) acknowledge the Ministry of Education, Government of India and IIT Mandi for the PhD fellowship. Nidhi Yadav (NY) acknowledges the Department of Biotechnology, Government of India, for the PhD fellowship. ASK and AS thank BioX centre, High-Performance Computing (HPC), and Advanced Material Research Centre (AMRC) at IIT Mandi for research facilities. ASK and NY are thankful to the Translational Health group of the International Centre for Genetic Engineering and Biotechnology (ICGEB), New Delhi, for providing research facilities and support. Biorender was used to prepare the schematics and the graphical abstract.

## Contributions

Study Conception and designing experiments-SKM, RKN, and ASK. Funding acquisition, resources, and supervision of the study-SKM and RKN. Data collection, validation, and analysis-ASK, AS, and NY. The first draft of the manuscript was written by ASK, SKM, and RKN. Data curation, formal analysis, and finalizing some of the experimental methodology-AS, NY. All authors contributed to reviewing and editing previous versions of the manuscript. All authors read and approved the final manuscript.

## Funding

This work was supported by the Core budget from the ICGEB, New Delhi, to RKN, and Seed grant from IIT Mandi to SKM.

## Declarations

### Ethical approval

The study is compiled with ethical standards.

### Competing interests

The authors declare no competing interests.

### Data availability

The RNA-sequencing data generated in the current study have been deposited in the NCBI Sequence Read Archive (SRA) under the BioProject accession PRJNA1517992.

## Notes

https://www.ncbi.nlm.nih.gov/

### Competing Interest Statement

The authors have declared no competing interest.

